# A pan-cancer analysis of the oncogenic role of COVID-19 risk gene Leucine Zipper Transcription Factor-Like Protein 1 (LZTFL1) in human tumors

**DOI:** 10.1101/2022.08.14.503890

**Authors:** Jihao Mo, Zhenzhen Zhang, Daping Wang, Mingqin Su, Jian Hu, Yakun Liu, Lei Wang, Meimei Wang

## Abstract

Population-based studies showed that COVID-19 infection causes higher death rate in cancer patients. However, the molecular mechanism of COVID-19 with cancer is still largely unknown. Here we analyzed the Leucine Zipper Transcription Factor-Like Protein 1 (LZTFL1) which is the most significant gene associated with COVID-19. First, we explored the potential oncogenic roles of LZTFL1 through transcriptome data from The Cancer Genome Atlas (TCGA) and Gene Expression Omnibus (GEO) database. LZTFL1 is significantly low expressed in 11 of 34 kinds of cancers we analyzed. Consistent with the mRNA expression data, the protein expression of LZTFL1 in lung adenocarcinoma (LUAD), clear cell renal cell carcinoma (ccRCC), Uterine corpus endometrial carcinoma (UCEC), and ovarian cancer (OV) patients are significantly decreased compared to healthy tissues. The survival analysis from the Kidney renal clear cell carcinoma (KIRC), Rectum adenocarcinoma (READ), and Uveal Melanoma (UVM), the LZTFL1 high expression group have a significantly higher survival rate compared to the low expression group. Taken together, LZTFL1 acts as a cancer suppressor gene for several cancers. Moreover, LZTFL1 expression was associated with the cancer-associated fibroblast infiltration in several tumors including Bladder Urothelial Carcinoma (BLCA), Breast invasive carcinoma (BRCA), Esophageal carcinoma (ESCA), Head and Neck squamous cell carcinoma (HNSC), Lung squamous cell carcinoma (LUSC), and Pancreatic adenocarcinoma (PAAD). Gene ontology analysis showed that cilium organization, positive regulation of establishment of protein localization to telomere and SRP-dependent cotranslational protein targeting to the membrane were involved in the function mechanisms related to LZTFL1. Our studies offer a relatively comprehensive understanding of the oncogenic roles of LZTFL1 across different kinds of tumors.

## Introduction

As of Oct 11, 2022, 622 million COVID-19 infection cases and 6.56 million deaths reported worldwide. The SPIKE protein of COVID-19 binds to the ACE2, then cleavaged by TMPRSS2 to infect the mammalian respiratory cells[1]. The new discovered COVID-19 target protein named Neuropilin 1, might handing the key to a new door for the virus to replicate in the human airway cells[2]. A survey published by the American Cancer Society Cancer Action Network showed that COVID-19 pandemic challenges more to cancer patients and survivors, including delays and cancellations of health care services, economic challenges affecting their ability to pay and so on[3]. Recent publications showed that the immunosuppressed status of some cancer patients increases their risk of infection and result in serious complications compared with the general population[4]. The genomics mutations related to cancer are well-studied compared to other diseases. However, how the genetic mutations related to COVID-19 is still largely unknown.

Genome wide Association Study of Severe and critical illness in COVID-19 patient samples indicated that Leucine Zipper Transcription Factor-Like Protein 1 (LZTFL1), tyrosine kinase 2 (TYK2), dipeptidyl peptidase 9 (DPP9), Interferon Alpha and Beta Receptor Subunit 2 (IFNAR2), C-C Motif Chemokine Receptor 2 (CCR2), and Alpha 1-3-N-Acetylgalactosaminyltransferase and Alpha 1-3-Galactosyltransferase (ABO) genes are closely associated with severe COVID-19[5, 6]. There is an urgent need to investigate the role of these genomics mutations in cancer. Among all these genes, LZTFL1 is the most significant gene associated with COVID-19. LZTFL1 is known to interact with Bardet-Biedl Syndrome (BBS) proteins[7]. Recent evidence has shown that LZTFL1 was a novel cancer suppressor gene associated with certain kinds of cancers[8]. Immunohistochemical (IHC) staining on 311 paired normal/cancer tissue arrays suggested that LZTFL1 suppresses gastric cancer cell migration[9]. Evidence from breast cancer patients indicated that microRNA-21 promotes breast cancer proliferation and metastasis by suppressing LZTFL1[10]. Besides, LZTFL1 suppresses lung tumorigenesis by maintaining differentiation of lung epithelial cells[11]. However, there is still no pan-cancer evidence on the relationship between LZTFL1 and all the tumour types based on bid clinical data analysis.

Here we conduct a pan-cancer analysis of COVID-19 associated gene LZTFL1 using TCGA and GEO databases. We investigated the potential molecular mechanism of LZTFL1 in the pathogenesis and clinical prognosis of different cancers by analysing the gene expression, survival status, DNA methylation, genetic alteration, and relevant cellular pathway. To study the function of LZTFL1 as a transcription factor, we predicted the top 100 downstream genes of LZTFL1 and analysed their expression in COVID-19 related RNA-sequencing data set. Furthermore, we analysed the function, localization, and microRNA target of LZTFL1 downstream genes.

## Materials and methods

### Genetic alteration analysis

The genetic alteration characteristics of LZTFL1 in the TCGA database were analysed using the cBioPortal (https://www.cbioportal.org/)[12]. Briefly, “LZTFL1” was entered for queries of genetic alteration characteristics. The mutated sites were labelled on the localization on the amino acid position on protein coding region with several mutations observed. The alteration frequency from TCGA data set was calculated with mutation, amplification, and deep deletion information.

### Structure prediction

The structure of LZTFL1 was predicted based on the NCBI Reference sequence NP_065080.1 using SWISS-MODEL (https://swissmodel.expasy.org/)[13]. The structure of amino acid sequence 1-64 was built based on template SMTL ID: 5wo3.1[14]. The structure of amino acid sequence 148-222 was built based on template SMTL ID: 3o0z.1[15].

### mRNA expression analysis

The systematical analysis of LZTFL1 mRNA expressions between tumour and adjacent normal tissues in TCGA database were conducted using “Gene_DE” module of tumor immune estimation resource, version 2 (TIMER2, http://timer.cistrome.org/)[16]. The box plot with tumour and adjacent normal tissues was labeled gray. The statistical significance computed by the Wilcoxon test is annotated by the number of stars (*: p-value < 0.05; **: p-value < 0.01; ***: p-value < 0.001). The LZTFL1 expression in single cell RNA-seq data was obtained from the study of Allergic inflammatory memory in human respiratory epithelial progenitor cells using Single cell portal [17].

### Protein expression analysis

The systematical analysis of LZTFL1 expressions between tumour and adjacent normal tissues in TCGA database were conducted using Clinical Proteomic Tumour Analysis Consortium (CPTAC) of UALCAN portal (http://ualcan.path.uab.edu/analysis-prot.html)[18]. The LZTFL1 expression of breast cancer, ovarian cancer, Colon cancer, Clear cell renal cell carcinoma, Uterine corpus endometrial carcinoma, and lung adenocarcinoma tumor and normal tissues were analyzed, respectively. The expression of LZTFL1 in each individual cancer stages cancer type with further investigated.

### Survival analysis

The survival analysis of LZTFL1 between high and low expression groups in TCGA database was conducted using “survival map” module of GEPIA2 (http://gepia2.cancer-pku.cn/#survival)[19]. The “overall survival” was used as the method. The group cutoff was set as the median. The cutoff-high and cutoff-low were set as 50%. The log-rank test with a 95% confidence interval was used in the analysis of hypothesis test.

### Immune infiltration analysis

The Immune infiltration analysis of LZTFL1 in EPIC, MCPCOUNTER, XCELL, and TIDE database was conducted using “Immune_Gene” module of TIMER2[16].The cancer-associated fibroblasts were selected for analysis. The P value and correlation value were calculated by Spearman’s rank after purity adjustment. The heatmap and scatter plot were used to visualize the data.

### Gene ontology analysis

The proteins which interacted with LZTFL1 in *homo sapiens* were investigated using the STRING tool (https://string-db.org/)[20]. The edges indicate both functional and physical protein associations and the line thickness indicates the strength of data support. The number of interactors to show for the first and second shell were set as no more than 20. The top 100 genes that interact with LZTFL1 based on co-expression were analysed with Harmonizome (https://maayanlab.cloud/Harmonizome/)[21]. The list of genes was further analysed in COVID-19 related GEO data sets using the Enrichr tool (https://maayanlab.cloud/Enrichr/)[22]. The biological function, subcellular localization, and miRNA regulation of LZTFL1 were visualized using a heatmap.

## Results

### Genetic alteration of LZTFL1 in cancer

In this study, we analysed the mutation site of LZTFL1 in the amino acid sequence. As shown in Fig. 1A, there are totally of 41 mutations observed in the LZTFL1 coding region. The most observed mutation point is R19. R19C was detected in rectal adenocarcinoma (READ) patient and R19H were detected in uterine endometrioid carcinoma (UCEC) or mucinous adenocarcinoma of the colon and rectum (COAD) patient based on the TCGA database. Other mutations observed near this location is R22G/H, K24N, and R28K/S. The Leucine Zipper (Leu_zip) domain locates at 20-294 amino acids. These mutations located at 19, 22, 24, and 28 are the starting point of the Leu_zip domain. Another location with multiple mutations accumulated at the binding region of the zipper structure. These mutations are E148K, E160D, E162D, and E172D/K. All the mutations above shortened the length of side chain. The most possible explanation is that these mutations altered the formation of homo dimer of the leucine zipper in the beginning and the closely localized zipper position, further affected the function of LZTFL1 as a transcription factor (Fig 1B&C).

**Figure 1.**
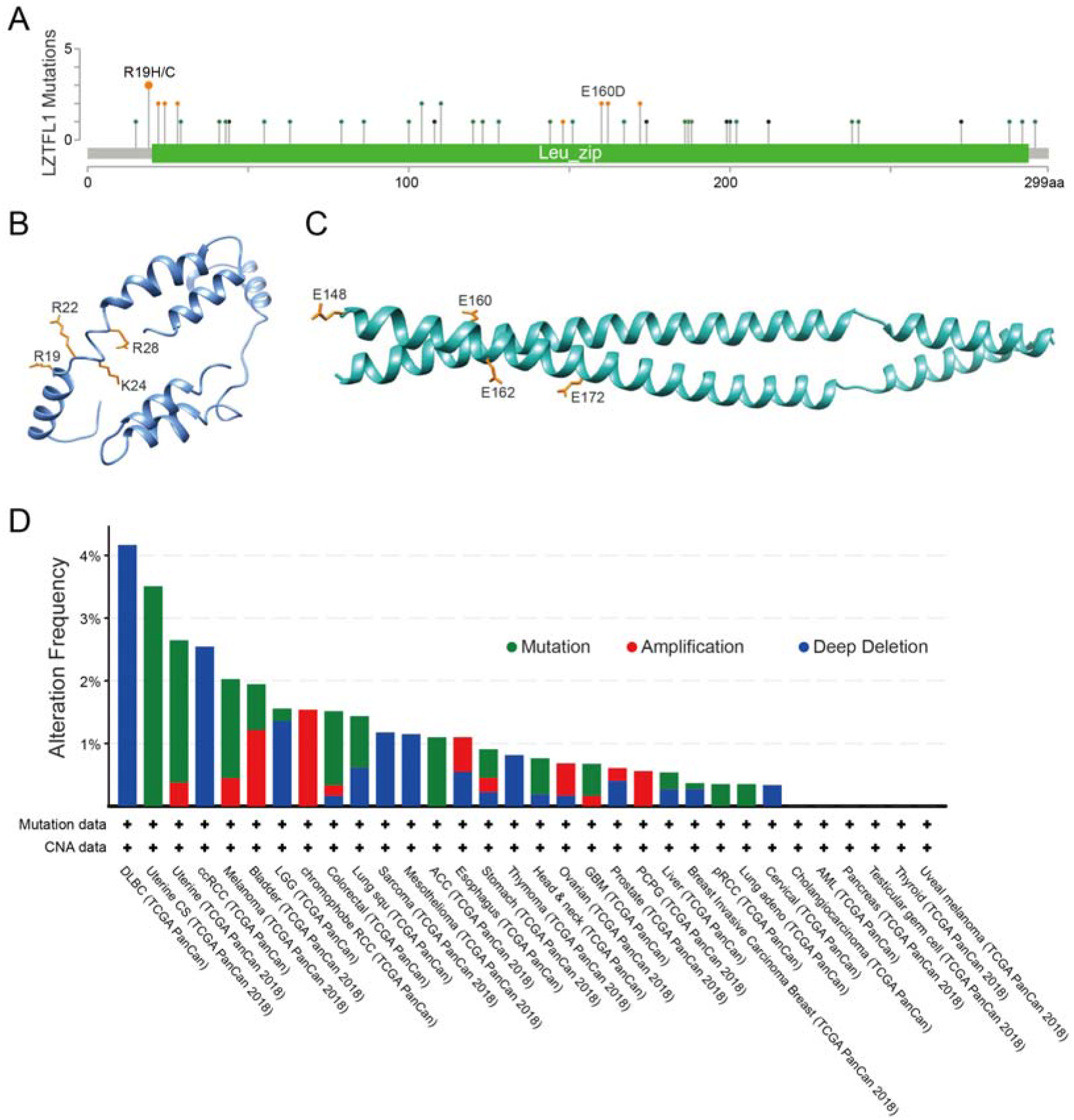
Mutation feature of LZTFL1 in different tumors. (A) The mutation site of LZTFL1 in tumors. The prediction of Leucine Zipper (Leu_zip) domain (B) and the binding region (C) structure of LZTFL1. The mutation type of LZTFL1 in different tumors.

We further analysed the genetic alteration of LZTFL1 in different tumour samples from TCGA database. Most of the LZTFL1 mutations are deep deletions. The highest alteration frequency of LZTFL1 (around 4%) appears for lymphoid neoplasm diffuse large B-cell lymphoma (DLBC) patients (Fig 1D). Only deep deletions are found in DLBC, ccRCC (clear cell renal cell carcinoma), sarcoma, mesothelioma, thymoma, and cervical patients. Only mutations are found uterine, adenoid cystic carcinoma (ACC), papillary renal cell carcinoma (pRCC), and lung adeno patients. Only amplification found in coulrophobe renal cell carcinoma (RCC) and pheochromocytoma/paraganglioma (PCPG) patients.

### Gene and protein expression of LZTFL1 in cancer

We further analyzed the expression of LZTFL1 in different types of cancers and paired non-tumor tissues in TCGA database. As shown in Fig.2A, LZTFL1 shows significantly increased expression in cholangiocarcinoma (CHOL), esophageal carcinoma (ESCA), head and neck squamous cell carcinoma (HNSC)-HPV+, and STAD (stomach adenocarcinoma) tumor groups. However, LZTFL1 shows significantly decreased expression in breast invasive carcinoma (BRCA), cervical squamous cell carcinoma and endocervical adenocarcinoma (CESC), colon adenocarcinoma (COAD), kidney Chromophobe (KICH), kidney renal clear cell carcinoma (KIRC), kidney renal papillary cell carcinoma (KIRP), liver hepatocellular carcinoma (LIHC), lung adenocarcinoma (LUAD), lung squamous cell carcinoma (LUSC), prostate adenocarcinoma (PRAD), skin cutaneous melanoma (SKCM), thyroid carcinoma (THCA), and UCEC (uterine Corpus Endometrial Carcinoma) tumor groups. To conclude, the mRNA expression of LZTFL1 plays different roles in different cancer types.

**Figure 2.**
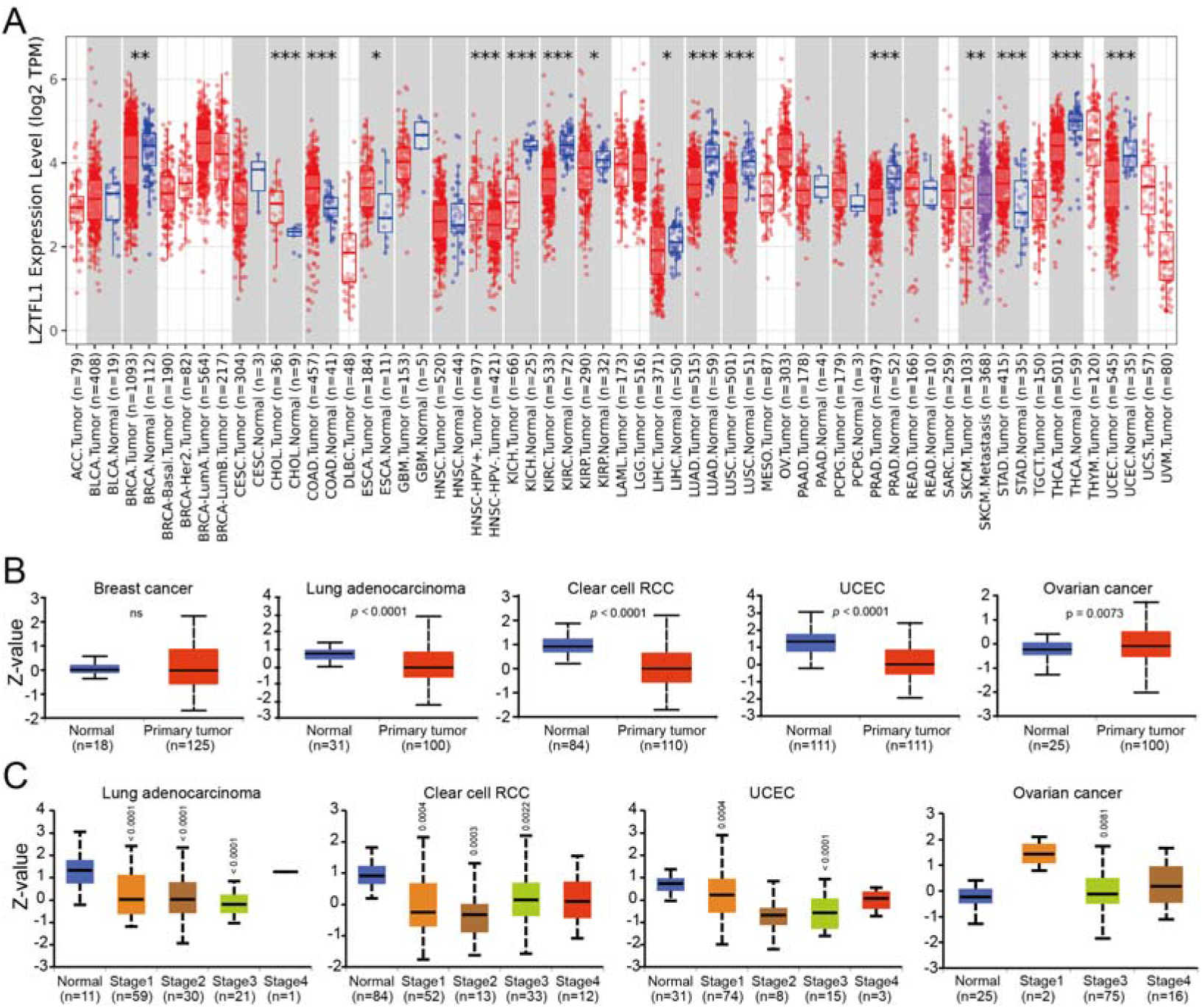
The mRNA and protein expression of LZTFL1 in different tumors and pathological stages. (A) The mRNA expression of LZTFL1 in different cancers or specific cancer subtypes was analyzed through TIMER2. (B&C) The protein expression of LZTFL1 in between normal tissue and primary tissue of breast cancer, lung adenocarcinoma, clear cell RCC, UCEC, and ovarian cancer by the main pathological stages.

We further analysed the protein expression data of LZTFL1 in CPTAC database. As shown in Fig 2B, the protein expression of LZTFL1 is significantly decreased in LUAD, Clear cell RCC, and UCEC patient samples. But the protein expression of LZTFL1 is significantly increased in ovarian cancer patient samples. To further investigate which stages of these cancers are correlated with LZTFL1 protein expression, we separated the data based on the stages. Patients with LUAD and clear cell RCC have decreased LZTFL1 protein expression in stages 1-3. Patients with UCEC have decreased LZTFL1 protein expression in stages 1 and 3. Patients with ovarian cancer have increased LZTFL1 protein expression in stage 3. These data indicated that the LZTFL1 protein expression is differentially associated with the prognosis of patients with different cancer types.

### Survival analysis of LZTFL1 in cancer

To investigate the correlation between the expression of LZTFL1 and the prognosis of patients with different kinds of tumours, we analysed the survival by dividing the cancer cases into high and low-LZTFL1 expression groups. The datasets from TCGA were used in this analysis. Based on the log rank *P*-value, patients with high LZTFL1 expression were linked to a better prognosis of overall survival for cancers of KIRC (*P* < 0.0001), READ (*P* = 0.0039), and UVM (*P* = 0.0180). patients with low LZTFL1 expression were linked to a better prognosis for cancers of Brain Lower Grade Glioma (LGG, *P* = 0.0100) (Fig. 3).

**Figure 3.**
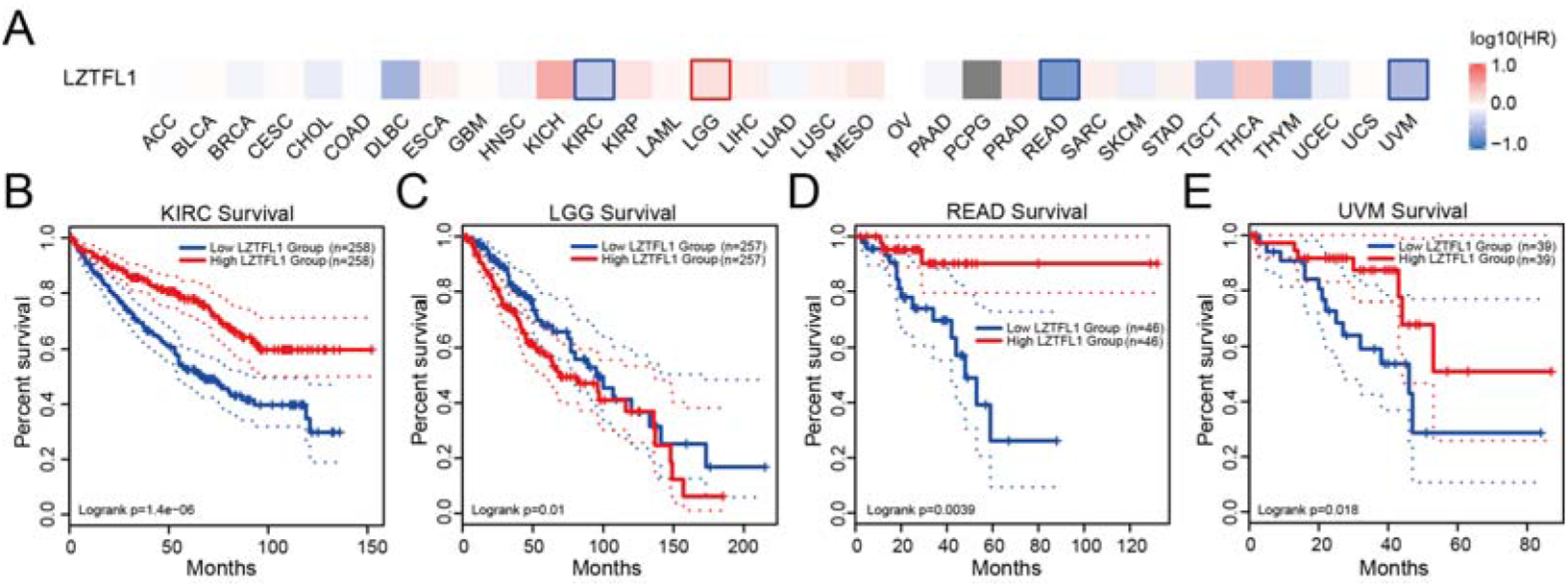
The correlation between LZTFL1 gene expression (A) and survival prognosis (B) of cancers in TCGA.

### Immune infiltration analysis of LZTFL1 in cancer

To analysis the immune infiltration of LZTFL1 across all kinds of tumours, the immune cells of cancer-associated fibroblasts were selected. The XCELL, MCPCOUNTER, TIDE and EPIC algorithms were applied to investigate the correlation between the cancer-associated fibroblasts and LZTFL1 gene expression. The statistical positive correlation of LZTFL1 expression and infiltration value of cancer-associated fibroblasts for the tumours of BLCA, BRCA, BRCA-basal, ESCA, HNSC, HNSC-HPV^-^, LUSC, and PAAD were observed (Fig. 4A). The scatterplot showed the related type of tumour using one algorithm (Fig. 4B).

**Figure 4.**
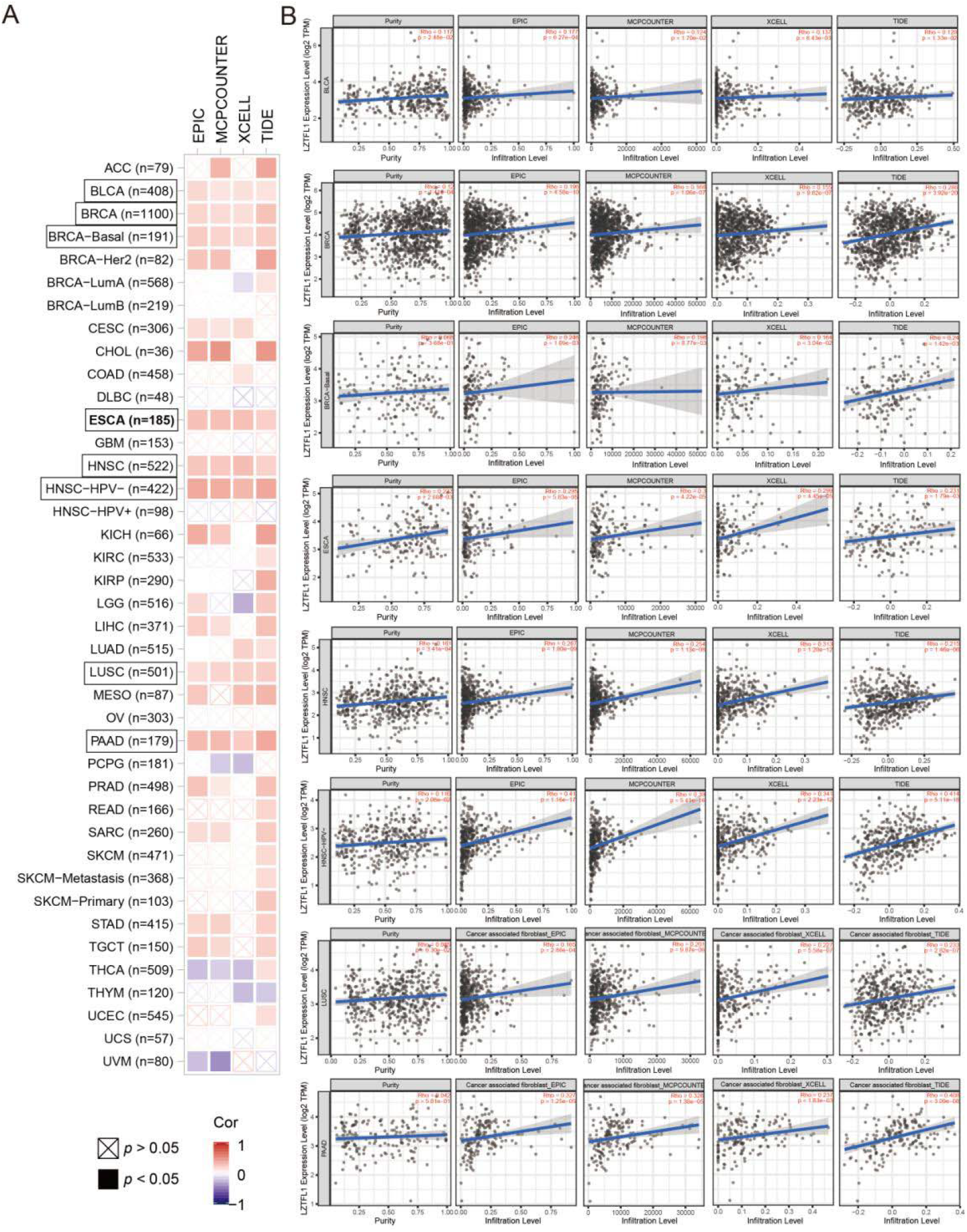
The correlation analysis between LZTFL1 expression and immune infiltration of cancer-associated fibroblasts. Different algorithms were used to explore the potential correlation between the expression level of the LZTFL1 gene and the infiltration level of cancer-associated fibroblasts across all types of cancer.

### Enrichment and downstream gene analysis of LZTFL1-related genes

To further investigate the molecular mechanism of the LZTFL1 gene in tumorigenesis, we analysed the LZTFL1 related genes by protein-protein interaction. Top 50 LZTFL1 related genes were enriched based on the co-localization, function, and co-expression analysis using STRING. The top 50 genes are enriched in cilium organization, SRP-dependent co-translational protein targeting to membrane, and positive regulation of establishment of protein localization to the telomere (Fig.5A). The target genes of LZTFL1 are found in transcriptome database of Down-regulated by SARS-CoV-2 in A549-ACE2 treated cells, Down-regulated genes from COVID-19 infected bronchoalveolar lavage from patients, MERS-CoV 38 Literature-Associated Genes from Geneshot, SARS perturbation Down Genes PBMCs, Down-regulated by SARS-CoV-2 infection in Vero E6, SARS Perturbation Down Genes Mouse Lung, SARS coronavirus endoRNAse from Virus-Host, MERS-Cov 6 Predicted Kinases Genes, Down Genes from SARS infected Mouse Lung, and SARS-CoV perturbation Down Genes Vero E6 (Fig.5B). The enriched pathways of LZTFL1 downstream genes are Collecting duct acid secretion, Vibrio cholerae infection, Synaptic vesicle cycle, Epithelial cell signalling in Helicobacter pylori infection, Vitamin digestion and absorption, Rheumatoid arthritis, RNA degradation, Oxidative phosphorylation, Sphingolipid metabolism, and mTOR signalling pathway (Fig.5C). The enriched cellular organelles of LZTFL1 downstream genes are Golgi trans cisterna, endosome membrane, filopodium, lysosomal membrane, tertiary granule membrane, integral component of ER, specific granule membrane, actin cytoskeleton, centrosome, and integral component of Golgi. The top 10 enriched miRNAs are hsa-miR-5683, hsa-miR-519d-3p, hsa-miR-20b-5p, hsa-miR-758-5p, hsa-miR-526b-3p, hsa-miR-7852-3p, hsa-miR-106a-5p, hsa-miR-17-5p, hsa-miR-20a-5p, and hsa-miR-4640-3p (Fig.5D&E).

**Figure 5.**
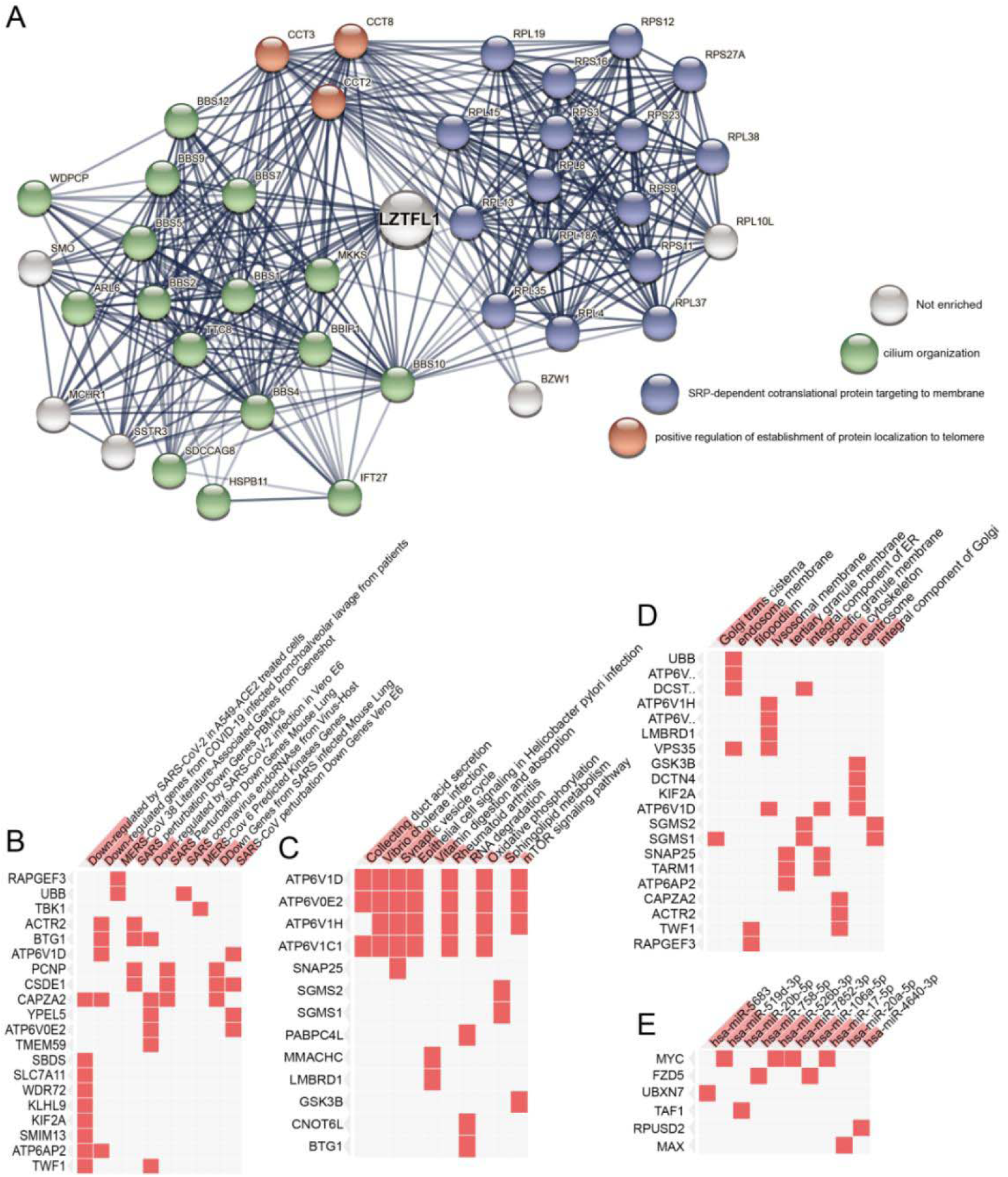
The LZTFL1 related enrichment and downstream gene function analysis. (A) The LZTFL1 related enrichment analysis. The downstream of LZTFL1 genes enriched in SARS-CoV-2 related GEO dataset (B) and human metabolism pathway (C). (D) The subcellular localization of LZTFL1 downstream gene. (E) The top 10 downstream miRNA regulated by LZTFL1.

## Discussion

LZTFL1 have been known as Bardet-Biedl Syndrome (BBS) proteins and smoothened trafficking regulator for around ten years[7]. LZTFL1 interacts with a BBS protein complex and regulates ciliary trafficking of this complex. A homozygous 5 bp deletion in LZTFL1 was identified in BBS patients, which causes no LZTFL1 transcript or protein could be detected in the patient’s fibroblasts[7]. LZTFL1 locals in cytoplasm helping with the regulation of BBS protein complex ciliary trafficking.

How LZTFL1 works in each kind of cancer types are still largely unknown. The current hypothesis suggested that by interacting with E-cadherin and the actin cytoskeleton, LZTFL1 regulating the transition of epithelial cells to mesenchymal cells[24]. Moreover, Wang et al reported that LZTFL1 suppresses gastric cancer cell migration and invasion through regulating nuclear translocation of β-catenin[9]. By compare the protein expression level with normal tissues by immunohistochemistry, Further studies showed that overexpress LZTFL1 inhibits anchorage-independent cell growth *in vivo* and *in vitro*[11]. Our studies showed that LZTFL1 protein expression was closely related to lung adenocarcinoma, clear cell RCC, UCEC, and ovarian cancer. However, different from the ICH results, LZTFL1 protein expression was not changed in lung cancer patients, but significantly increased in ovarian cancer patients’ samples compare to normal tissues. Since the Qun et al’s studies use epithelial cell of ovarian from 10 patients and lung epithelial cell samples from 9 patients, the CPTAC database have 125 samples from breast and 100 samples from whole ovarian cancer samples, the different results might come from the difference of cell type and sample numbers.

As a transcription factor, the binding region of LZTFL1 is vitally important for regulating downstream gene expression. Mutations in the binding region of LZTFL1 have been reported related to different cancer types as well as COVID-19. Recent studies investigated that the rs17713054 risk allele of LZTFL1 generates a CCAAT/enhancer binding protein beta motif[24]. SNP allele of LZTFL1, rs17713054G>A, regulates the expression of LZTFL1 in lung samples by interacting with the LZTFL1 promoter. Further studies from lung samples of COVID-19 patients showed that a signal of epithelia-mesenchymal transition, which is essential for the innate immune response in lung epithelial cells[24]. Single cell plot of human respiratory epithelial progenitor cells showed that LZTFL1 were mainly expressed in cliated cells, which was consistent with the protein function analysis that LZTFL1 was required for cilium organization (Fig.6A). These findings are consistent with our results that most of the downstream genes regulated by LZTFL1 were down regulated in SARS-CoV-2 infected lung cells, Vero cells, mouse lung samples, and bronchoalveolar lavage samples from COVID-19 patients. Our hypothesis is that LZTFL1 serves as a protective role against the COVID-19 infection as well as the survival of KIRC, READ, UVM cancer patients (Fig.6B).

**Figure 6.**
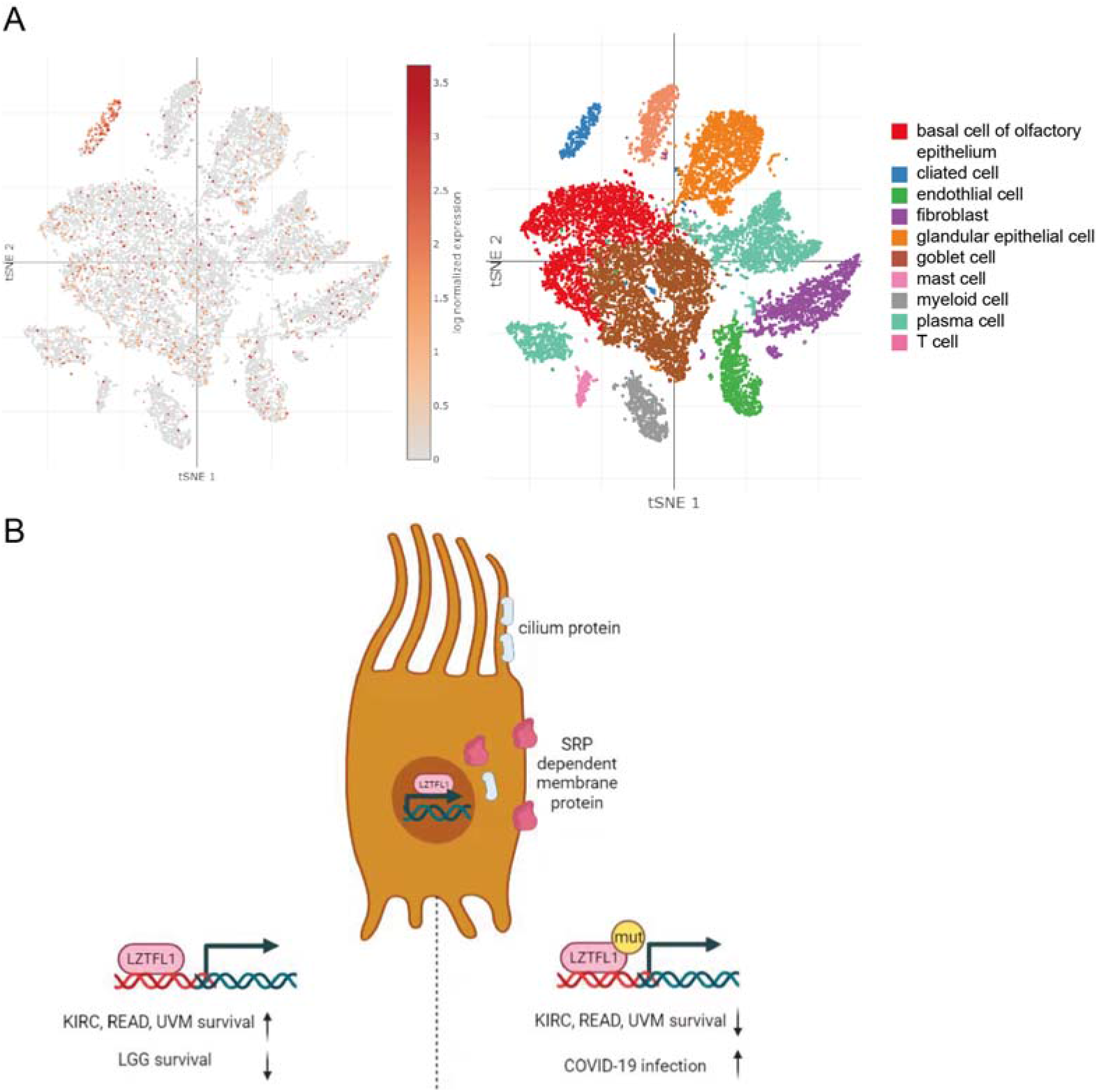
(A)Single cell plot of LZTFL1 expression in human respiratory epithelial progenitor cells. (B)The schematic diagram shows the role of LZTFL1 in COVID-19 and cancer.

## Acknowledgments

This work was funded by Project supported in part by the National Science Fund for Distinguished Young Scholars of China (No. 82201315), Anhui Provincial Natural Science Foundation (2008085MB49) and Funded Project of Anhui Medical University’s Research Level Improvement Program (2021xkjT004).

